# Adaptive sequential eye-movement sampling and replay under different task demands

**DOI:** 10.64898/2026.02.16.706087

**Authors:** Qiaoli Huang, Christian F. Doeller

## Abstract

Human cognition is capacity-limited, requiring strategies to actively structure information. Eye movements offer a natural mechanism for sequential sampling, but whether such sequences organize mnemonic representations is unknown. We developed a working-memory task where color-frequency pairings created a consistent latent ordinal structure to optimally reduce memory load. Across two experiments, gaze patterns spontaneously aligned with this structure.

Participants sampled items following this sequence during encoding and covertly replayed them during maintenance. Critically, the expression of this structure depended on cognitive demand. In a 3-item task, high performers showed robust sequential sampling during encoding, whereas lower performers compensated with replay-like revisitation during maintenance. Under higher demand (4 items), encoding-based organization was disrupted, and structured replay emerged primarily during maintenance to support memory. These findings show that eye movements do more than reflect memory; they actively organize it, revealing a flexible, behavioral analogue to neural replay when encoding resources are strained.

## Introduction

The human mind operates under strict capacity limits: we cannot process or store every detail of the complex visual world. To optimally deal with these constraints, perception and memory have evolved to rely on efficient coding strategies that exploit the statistical regularities of the environment (Attneave, 1954; Ganguli & Simoncelli, 2014; Wei & Stocker, 2015). Real-world scenes are generally structured and predictable, allowing us to use prior knowledge to guide goal-directed behavior. For example, when searching for a book in a library, you do not scan shelves at random; instead, you use the numerical or alphabetical order of catalog labels to predict where the book should be and then move sequentially along the shelves to find it efficiently. Such strategies depend on orderly sampling of structured information, a process that, in vision, is carried out through eye movements.

Because the central fovea offers high acuity but only over a very limited region of the visual field, our eyes must continually move to sample different parts of a scene. This active sensing strategy not only enables efficient visual processing but also provides a unique window into memory systems (Hannula et al., 2010). Indeed, the oculomotor system is tightly integrated with the brain’s memory circuitry (Meister & Buffalo, 2016), with growing evidence demonstrating that eye movements actively shape memory formation and retrieval. As a result, gaze behavior can serve as a readout of both the accessibility and internal organization of memory contents (Althoff & Cohen, 1999; Hannula & Ranganath, 2009; Loetscher et al., 2010; Van Ede et al., 2019; Viganò et al., 2024; Zonca et al., 2020), often quantified through summary measures such as the number of fixations or the distribution of saccade vectors.

Building on this perspective, recent research has increasingly focused on the temporal organization of gaze, revealing that eye movements unfold as structured sequences rather than a series of independent fixations. In visually guided tasks, eye-movement trajectories often follow near-optimal policies for maximizing information gain (Hoppe & Rothkopf, 2019; Yang et al., 2016; Zhu et al., 2022). Such sequential sampling provides the visual system with an ordered stream of inputs, which may be especially valuable for constructing stable working-memory representations under capacity constraints. This view aligns with proposals that perceptual and mnemonic systems share efficient-coding principles (Bays et al., 2024), suggesting that the structure of eye-movement sequences may play an active organizational role in memory formation.

Yet despite these insights, the functional contribution of eye-movement sequences in mnemonic organization remains unclear. Two questions are particularly critical. First, how are sequential sampling orders established when multiple items must be encoded simultaneously—does the oculomotor system actively impose structure on concurrent inputs? Second, and more importantly, does this sequence persist or re-emerge during maintenance, when no visual input is available? Demonstrating such internal “replay” of gaze sequences would provide direct evidence that eye movements do not merely accompany memory but participate in its organization, paralleling neural replay phenomena thought to support consolidation and mental reorganization (Foster & Wilson, 2006; Huang et al., 2018; Liu et al., 2019; Liu, Mattar, et al., 2021; Schuck & Niv, 2019).

To address these questions, we developed a working memory paradigm that embeds a latent structure within the to-be-memorized items. Participants memorized multiple gratings presented simultaneously at distinct spatial locations, each varying in both color (green–blue continuum) and spatial frequency. This dual-feature design imposed substantial working memory demands, encouraging the use of efficient sampling strategies. Critically, spatial frequency was systematically coupled with color hue to create an implicit, ordinal structure across feature space, which could be leveraged to reduce memory uncertainty. This design allowed us to test whether participants would adopt the corresponding structured sampling sequence during encoding and, moreover, whether this sequence would re-emerge during maintenance when stimuli were no longer visible. Finally, by varying memory load across two experiments, we assessed how cognitive demand modulates reliance on encoding-based versus maintenance-based gaze organization, thereby clarifying the functional role of oculomotor sequencing in supporting working memory.

## Results

### Eye-movement trajectory is modulated by the spatial configuration of the memory contents

Participants were briefly shown three gratings presented simultaneously at distinct spatial locations, each varying in spatial frequency and color. At the end of each trial, they reported the location of a probed feature (either spatial frequency or color) or indicated that the feature was new (Figure 1A). This task challenged working memory capacity by requiring the maintenance of multiple feature–location bindings. To probe potential efficient coding strategies, we introduced an implicit regularity in the stimulus set: lower-frequency gratings were consistently paired with “greener” hues. This contingency established an ordinal structure among the items, providing a scaffold to reduce memory load. Crucially, on every trial, items could be ranked by their relative position (value) on this feature gradient (item 1: lowest frequency/greenest; item 2: intermediate; item 3: highest frequency/least green). These ordinally labeled items were assigned to fixed locations, yielding six possible configuration patterns (Figure 1B). To encourage natural encoding strategies, participants were allowed to move their eyes freely except when a central fixation dot was presented, during which they were required to maintain fixation. Our analyses focused on eye-movement patterns during both the encoding period (when the memory items were visible) and the maintenance period (after their disappearance).

**Figure 1.**
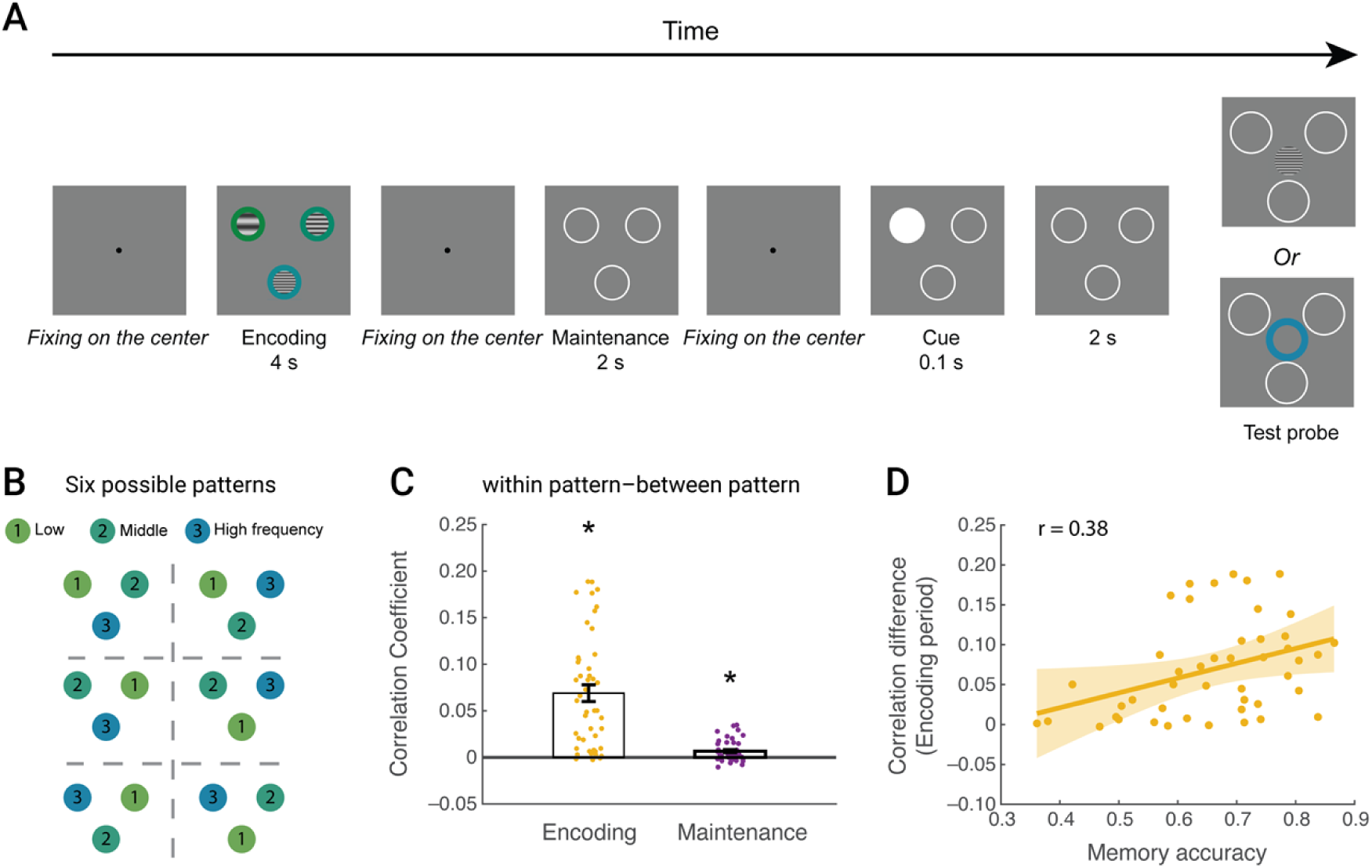
Experimental paradigm and structured gaze trajectory. **(A)** Experimental paradigm (N = 46). Participants began each trial by fixating on a central dot. Three gratings (varying in color and spatial frequency) then appeared simultaneously for 4 s, at three fixed, equidistant locations around the center (7° eccentricity). Participants were instructed to memorize each item’s color, frequency, and location. After the gratings disappeared, a central fixation dot reappeared, indicating participant to re-fixate. Next, three white circles occupying the same locations as the gratings were presented for 2 s, followed by another fixation period. Then, the three circles reappeared with one filled for 100 ms to direct attention to that specific location, followed by another 2 s display of all circles. In the end, a test stimulus (color or frequency) appeared at the center of the screen, and either matched one of the memorized items or was novel. Participants needed to report the associated location or indicate “new”. The cue was completely invalid because the test stimulus was equally likely to match the memory item at any given location. Participants can freely move their eyes except the fixation period. **(B)** Stimulus configuration patterns. There were six spatial configuration patterns based on the relative feature values. Item 1: lowest frequency, greenest; Item 2: middle frequency, intermediate hue; Item 3: highest frequency, least green. **(C)** Pattern-similarity effect. The contrast of the correlation (mean ± SEM) between within-pattern and between-pattern trial pairs during the encoding period (yellow) and maintenance periods (purple). Each dot indicates individual participants (*: p<0.05). **(D)** Relationship with memory performance. Scatterplot shows memory accuracy (X-axis) versus pattern similarity effect in the encoding period (Y-axis).

Behavioral performance was well above chance level (0.25) across all conditions (one sample t-test: overall t _(45)_ = 22.46, p < 0.001, Cohen’s d = 3.31; color domain t _(45)_ = 22.87, p < 0.001, Cohen’s d = 3.37; frequency domain t _(45)_ = 15.50, p < 0.001, Cohen’s d = 2.29). We then asked whether the embedded cross-feature ordering structure actively guided eye-movement trajectories. Specifically, if participants sampled information based on this latent item order, then eye-movement trajectories would be more similar for trials sharing the same configuration (within-pattern) than for trials with different configurations (between-pattern). To test this, we quantified attentional allocation using gaze-to-item distance time courses (Figure 2A), where shorter distances reflected stronger engagement/representation with a given item. For each item, we computed pairwise correlations of these time courses across trials and categorized each pair as within-pattern or between-pattern depending on whether the trials shared the same configuration or not, and then averaged the correlations across items to yield a single summary measure per participant (see Methods). The contrast between within– and between-pattern correlations quantified the pattern-similarity effect, providing an index of value-guided sampling while controlling for any baseline biases associated with spatial location.

**Figure 2.**
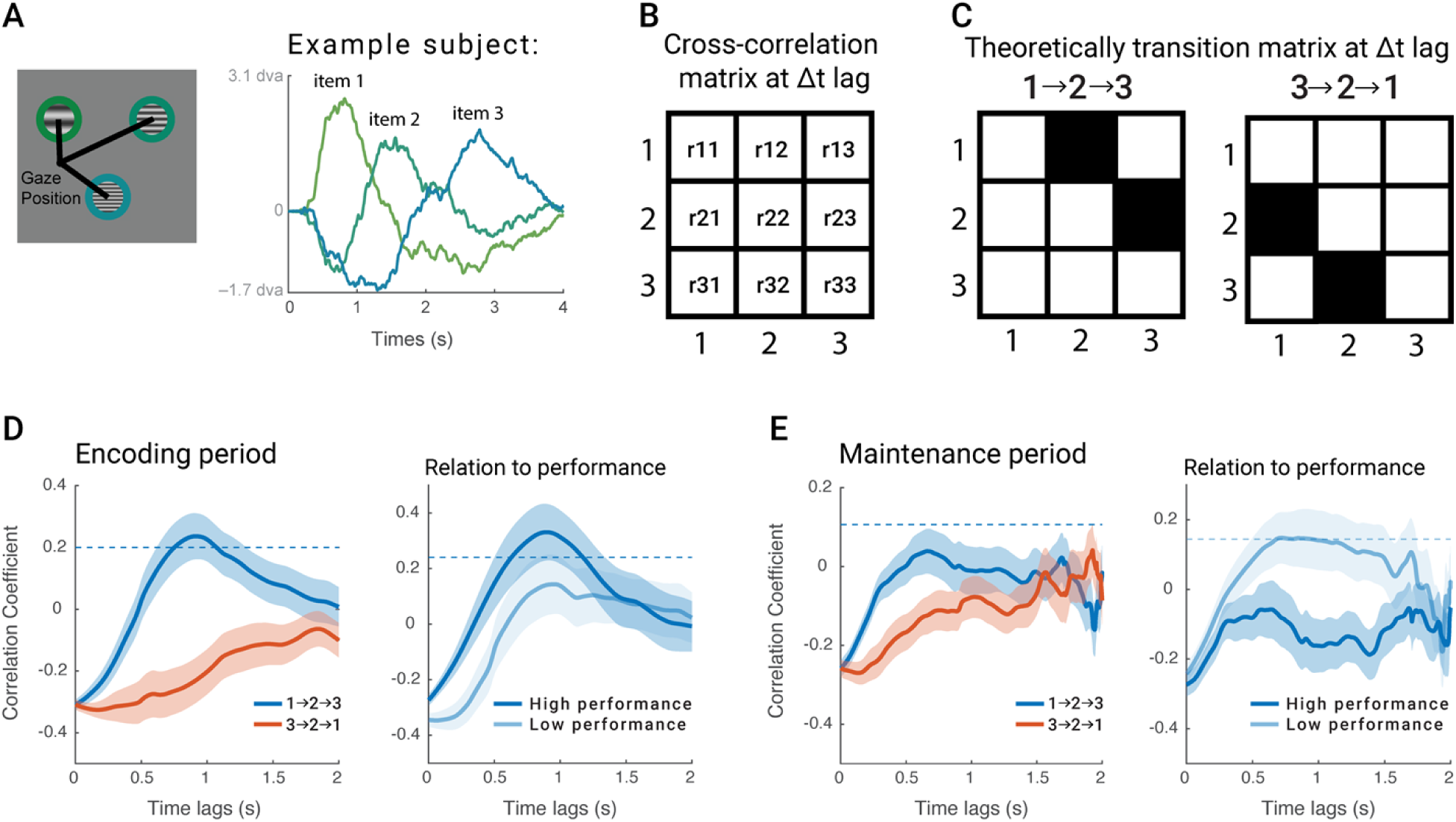
Sequential eye-movement sampling and replay. **(A)** Gaze-to-item distance analysis. Left: schematic illustrating the calculation of the gaze distance to each item. Right: time course of gaze distances to individual items (averaged across trials) for a representative participant, with items labeled by their relative values. **(B)** Empirical gaze cross-correlation. The matrix depicts the temporal relationship between gaze patterns for each item at a specified time lag (Δt). **(C)** Theoretically transition model: 1→2→3 sequence (left panel); 3→2→1 sequence (right panel). **(D)** Sequence strength during encoding. Left: strength of 1→2→3 (blue) and 3→2→1 (red) sequences, separately quantified by the correlation between empirical cross-correlation matrix (B) and transition models (C). Right: 1→2→3 sequence strength in high (N = 23, dark blue) and low-performance (N = 23, light blue) groups. The dashed line indicates the statistical threshold (p < 0.05) derived from permutation testing, corrected for multiple comparisons. **(E)** Sequence strength during maintenance. Sequence strength analysis, equivalent to (D), was performed during the maintenance period of the task.

As predicted, we observed a significant pattern effect, i.e., gaze trajectories were more similar within-pattern than between-pattern during both the encoding (t_(45)_ = 7.74, p < 0.001, Cohen’s d = 1.14) and maintenance period (t_(45)_ = 4.13, p < 0.001, Cohen’s d = 0.61, Figure 1C). This effect remained when we balanced trial numbers across within– and between-pairs (see supplementary figure 1A). Critically, participants who showed a stronger pattern-similarity effect during encoding also exhibited higher memory accuracy (r = 0.38, p = 0.01, Figure 1D). This suggests that value-guided gaze trajectory during encoding reflects more structured mnemonic processing, which in turn leads to better memory performance.

### Sequential eye movements reveal performance-dependent sampling and replay

Having established that gaze patterns reflect structured, value-guided mnemonic processing, we next examined the temporal organization of these trajectories. Specifically, we asked whether participants sampled items in a systematic sequence that followed the shortest “mental path” defined by their relative frequency values (either 1→2→3 or 3→2→1, where 1–3 correspond to items ordered from lowest to highest spatial frequency). To quantify sequential structure, we applied a sequenceness-analysis framework (Liu, Dolan, et al., 2021). We first computed cross-correlation coefficients between all item pairs to generate a correlation matrix for each time lag (Figure 2B). Next, we specifically tested two transition models against these empirical matrices: a 1→2→3 model (where item 1 at time T predicts item 2 at T+Δt, and item 2 at time T predicts item 3 at T+Δt) and a reverse 3→2→1 model (Figure 2C). Permutation tests with item-label shuffling (see Methods) showed that only the 1→2→3 sequence significantly explained the empirical data, with a prominent effect at lags of 0.75–1.07 s (peak = 0.91 s, Figure 2D, left). To assess behavioral relevance, we split participants into high– and low-performance groups based on a median split of accuracy (N = 23 per group). Strikingly, the high-performance group exhibited a robust 1→2→3 sequential profile (lags: 0.63–1.17 s; peak = 0.89 s), whereas the low-performance group did not (Figure 2D, right). These results indicate that this ordered sampling strategy supports more effective memory encoding.

Having characterized the sequential structure of encoding, we turned to the maintenance period to determine whether this sequence was replayed during the maintenance period to potentially stabilize or consolidate structured mnemonic representations. To address this, we applied the same analysis to the maintenance period. At the group level, no significant sequential eye-movement pattern was observed, suggesting that explicit overt “replay” is not a general maintenance mechanism in this task. However, analysis of individual differences revealed a striking and unexpected dissociation: contrary to what might be assumed, the low-performance group exhibited significant sequential “re-visitation” at 0.68 – 0.95 s lags (peak: 0.72 s), whereas the high-performance group showed no evidence. Importantly, control analyses confirmed this effect was not explained by spatial biases, as sequence analysis based solely on item locations yield no significant structure (supplementary figure 2).

Taken together, these findings demonstrate that sequential eye-movement patterns constitute an effective strategy for structuring multiple items in memory, and that the strength of this structure reliably predicts performance. Critically, we observed a clear dissociation between encoding and maintenance periods as a function of individual ability. High-performing participants showed robust value-ordered sampling during encoding (lowest → highest) but no evidence of sequential replay during maintenance, consistent with efficient initial encoding that reduces the need for subsequent stabilization. In contrast, low-performing participants failed to deploy this encoding strategy and instead relied on replay during maintenance to reorganize and stabilize memory traces. This behavioral pattern aligns with evidence from neural replay, where replay primarily supports the consolidation of weakly encoded information, whereas strong initial encoding diminishes the need for later reactivation (Huelin Gorriz et al., 2023; Schapiro et al., 2018).

### Retro-cue initiates sequential recall following the order

To further validate the observed sequential organization observed in earlier analyses, we examined eye-movement sequence following a retro-cue, which is presented immediately after the maintenance period to exogenously direct attention and saccade. In principle, an effective cue should draw both attention and initial gaze to the cued item, after which subsequent saccades may reveal the internal priority structure of the remaining items. Given the observed 1→2→3 sequence during encoding/maintenance, we predicted that: (1) when item 1 was cued, the second saccade would more likely target item 2 than item 3, and (2) when item 2 was cued, it would favor item 3 over item 1. We first verified cue effectiveness by analyzing first-saccade destinations (Figure 3A). Fixation heatmaps following cue onset revealed a strong spatial bias toward the cued locations (Figure 3B). Quantitatively, first saccades landed near the cued location significantly more than chance (1/3; t _(45)_ = 4.30, p < 0.001, Cohen’s d = 0.63; Figure 3C).

**Figure 3.**
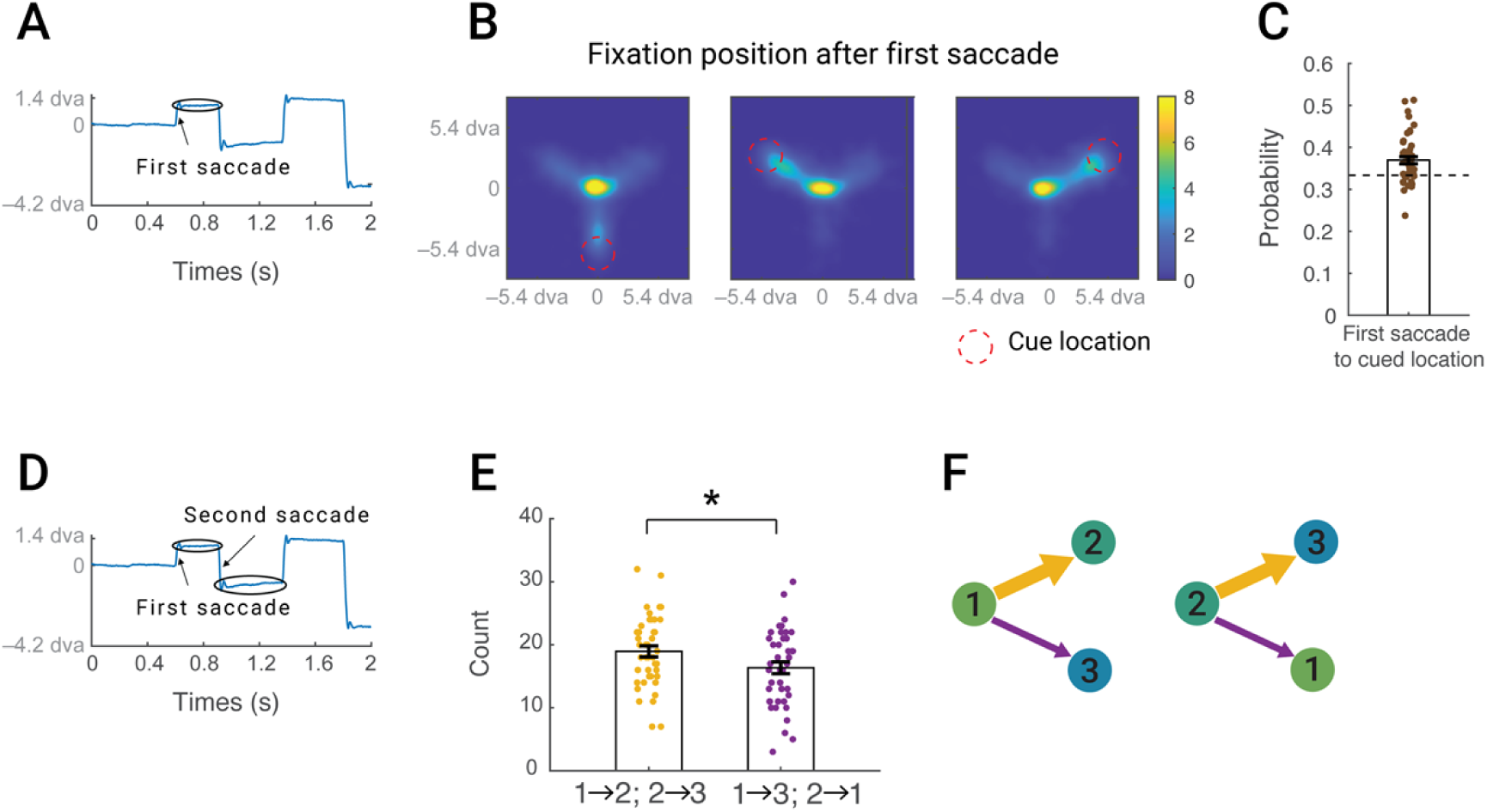
Ordered eye-movement transition after retro-cue. **(A)** Example gaze trajectory (y-axis position over time) illustrating the first saccade after retro-cue onset. **(B)** First-saccade destination heatmaps for each cue location (bottom-below, left-up, right-up). Red dashed circles mark the cued locations. **(C)** Proportion of the first saccade directed toward the cued location. Each dot indicates an individual participant. Dashed line indicates the chance level (1/3). **(D)** Example trajectory showing the second saccade following the retro-cue. **(E)** Second-saccade transition frequencies: left bar (mean ± SEM) shows predicted sequential transitions (1→2, 2→3); right bar (mean ± SEM) shows alternatives (1→3, 2→1). Each dot indicates an individual participant. **(F)** Schematic illustrating the 1→2→3 sequence. Preferred transitions (large arrows) occur more frequently than non-preferred transitions (small arrows): after recalling item 1, gaze is more likely to move to item 2 than item 3; after recalling item 2, it is more likely to move to item 3 than back to item 1.

To test whether the second saccade (Figure 3D) followed the predicted sequential order, we focused on trials that met three criteria: (1) the first saccade landed on the cued location, confirming effective cueing; (2) either item 1 or item 2 (the positions from which ordered transition are interpretable) was cued; and (3) at least two saccades were made. Note that 4 participants were excluded from further analyses because fewer than 20 trials satisfied these requirements. Consistent with our hypothesis, second-saccade transitions exhibited strong directionality: item 1→item 2 and item 2→item 3 transitions occurred significantly more frequently than alternative transitions (item 1→item 3 and item 2→item 1) (t_(41)_ = 3.11, p = 0.003, Cohen’s d = 0.48; Figure 3E), which robustly validated the optimal 1→2→3 sequential organization. Together, these results demonstrate that recall-phase eye movements precisely mirror the same ordinal structure observed during encoding/maintenance, confirming the functional role of active sequential organization in memory representation.

### High cognitive load forces a shift from encoding-period organization to maintenance-period replay

Our previous results showed that structured gaze sequences support memory, but their implementation depends on available cognitive resources. High performers, with resources to spare, immediately applied sequential sampling during encoding, whereas low performers appear to shift this organizational effort into the maintenance period, relying on sequential revisitation to reorganize weaker traces. This dissociation raised a key question: Does increasing cognitive load systematically disrupt encoding-phase organization and promote a compensatory shift of structured eye movements from encoding to maintenance? To address this, we designed a higher-load 4-item memory task (Figure 4A), analogous to the 3-item paradigm but with increased demands (Figure 4A). Participants performed well above chance (0.2; t _(39)_ = 22.33, p < 0.001, Cohen’s d = 3.53), though accuracy was lower than in the 3-item task (Figure 4B: two independent sample t-test, t _(84)_ = 2.58, p = 0.01, Cohen’s d = 0.60). Note that after accuracy was adjusted for chance level, this difference remained marginal (t _(84)_ = 1.62, p = 0.05 (one-tailed), Cohen’s d = 0.35), confirming that the 4-item task imposed a greater cognitive demand.

**Figure 4.**
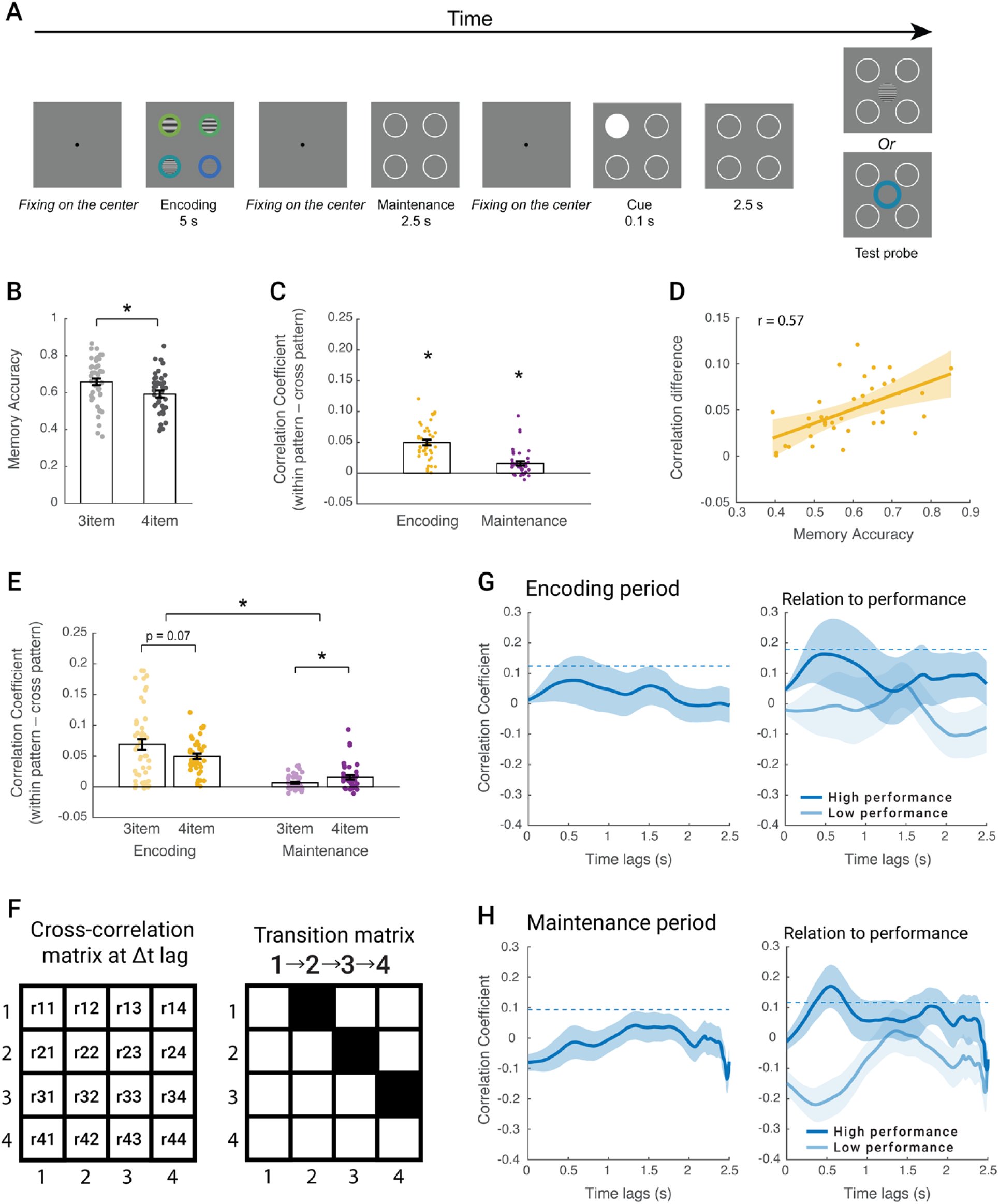
Experimental paradigm and results of 4-item task. **(A)** Experimental paradigm for the 4-item memory task (N = 40). The design was identical to the 3-item task, except that participants were asked to remember four items. **(B)** Comparison of memory accuracy (mean ± SEM) between 3-item (gray) and 4-item (black) tasks. Each dot indicates one participant. **(C)** Pattern-similarity effect. Difference in correlation (mean ± SEM) between within-pattern and cross-pattern trial pairs, shown separately for the encoding period (yellow) and maintenance period (purple). **(D)** Relationship between memory performance and pattern similarity. Scatterplot showing that participants with stronger pattern-similarity effects during encoding exhibited higher memory accuracy. **(E)** Comparison of pattern similarity effects (mean ± SEM) between 3-item (light color) and 4-item (dark color) tasks for both memory encoding (yellow) and maintenance (purple) periods. **(F)** Left: empirical correlation matrix at Δt lag formed by the pair-wise cross-correlation between items. Right: theoretically transition matrix corresponding to the 1→2→3→4 sequence. **(G**) Sequential organization during encoding. Left: time course of 1→2→3→4 sequence strength (mean ± SEM) across all participants during the encoding period. Right: sequence strength in high (N = 20, dark blue) and low-performance (N = 20, light blue) groups. Dashed lines indicate permutation-derived significance thresholds corrected for multiple comparisons. **(H)** Sequential organization during maintenance. Same as (G) but showing 1→2→3→4 sequence strength during the maintenance period.

Replicating our 3-item analysis, we first examined whether eye-movement trajectories also showed a pattern-similarity effect, i.e., greater similarity for trials sharing the same configuration pattern relative to trials with different patterns. This pattern effect was again significant during both encoding (t_(39)_ = 10.66, p<0.001, Cohen’s d = 1.69) and maintenance (t_(39)_ = 4.54, p < 0.001, Cohen’s d = 0.72, Figure 4C), with encoding effect strongly predicting memory accuracy (r = 0.57, p < 0.001, Figure 4D). Critically, a comparison between the 3-item and 4-item tasks revealed a significant interaction effect (Figure 4E, F_(1,84)_ = 6.84, p = 0.01, η_p_ ^2^ = 0.08): the 4-item task showed marginally reduced pattern effects during encoding (t _(84)_ = –1.84, p = 0.07, Cohen’s d = 0.40) but significantly stronger effect during maintenance (t _(84)_ = 2.43, p = 0.02, Cohen’s d = 0.53) compared to 3-item task. These findings suggest that as task demands increase, efficient organization during encoding becomes less feasible, prompting compensatory reorganizational processes during maintenance to support memory.

We next asked whether the more fine-grained sequential structure of eye movements showed a comparable shift. Following the analysis in the 3-item task, we constructed a 4 × 4 time-lagged cross-correlation matrices and test whether it reflected the 1→2→3→4 sequence (Figure 4F). During the encoding period, no significant group-level expression of this sequential pattern was observed (Figure 4G), although high-performing participants showed a non-significant trend toward structured sampling (see supplementary figure 3 for nonsignificant 4→3→2→1). In contrast, during the maintenance (Figure 4H), while the group again showed no significant sequence, high performers exhibited significant sequential re-visitation at lags of 0.35 – 0.74 s (peak at 0.55 s). This result resembled the maintenance-period re-visitation pattern observed in lower performers during the 3-item task, suggesting that under higher cognitive load, a compensatory, internally generated replay-like mechanism for structured representations.

Taken together, these two experiments demonstrate that eye movements actively support working memory by structuring information in an ordered sequence. Critically, the deployment of this organizational strategy depends on available cognitive resources. In the 3-item task, where efficient encoding is manageable, high performers engaged structured sampling during encoding, whereas low performers compensated by relying on sequential revisitation during maintenance. In the more demanding 4-item task, encoding-phase organization was disrupted for most participants, and only high performers preserved performance by developing structured replay during maintenance. Thus, under higher cognitive load, sequential reorganization during maintenance becomes essential for preserving information. More broadly, these findings indicate that sequential eye movements constitute an active, adaptive mechanism for internally structuring information in support of memory.

## Discussion

The fovea’s high acuity comes at the cost of limited spatial coverage, necessitating frequent eye movements to sample the visual input. Working memory faces a similar constraint: its limited capacity requires efficient strategies for organizing information during encoding and retention(Brady et al., 2011; Cowan, 2001). To investigate how eye movements contribute to such organization under complex environment, we asked participants to remember three simultaneously presented gratings that varied in color (from green to blue) and spatial frequency. This design largely challenged working memory capacity by requiring the maintenance of multiple feature–location bindings. Critically, lower spatial frequencies were consistently paired with “greener” hues. This association created a latent structure that afforded an optimal sampling sequence—either ascending or descending in feature space—that could help to reduce memory load. Across trials, eye-movement trajectories reflect structured mnemonic processing and directly predicted memory performance. High-performing participants expressed this structured sequence during encoding, whereas low-performing participants only reinstated it during maintenance, indicating a shift toward compensatory reorganization when initial encoding was weak. Under higher cognitive load, reliance on maintenance-period sequence replay increased further. Together, these findings show that eye movements do not merely accompany memory but reveal an internally generated organizational process that supports the retention of multiple items, especially when cognitive resources are strained.

Our key finding reveals that eye-movement strategies are dynamically reorganized depending on task demands. Under low demands, participants exhibited an optimal sequential sampling pattern during encoding. As demands increased,however, this structured strategy disappeared during encoding and re-emerged during the maintenance period, reflecting a reorganization of information after stimulus offset. This pattern parallels neural replay literature, where weakly encoded information elicits stronger offline reactivation (Huelin Gorriz et al., 2023; Schapiro et al., 2018), and aligns with eye-tracking evidence showing that stable or well-learned memories require less exploratory sampling (Althoff & Cohen, 1999; Gong et al., 2025). Together, these results suggest that when encoding demands exceed cognitive capacity, the brain compensates by developing optimal sequential resampling during the maintenance period (despite the absence of visual input) as an active mechanism for reorganizing and stabilizing fragile memory representations.

Our previous work using EEG/MEG decoding has shown that spontaneous neural replay occurs during working memory maintenance and retrieval, and that this replay supports behavioral performance (Huang & Doeller, 2026; Huang et al., 2018; Huang & Luo, 2024; Huang et al., 2021). Building on these findings, the present study uses eye movements as a high-resolution behavioral readout of replay-like processes and demonstrates that this reactivation is flexibly modulated by task demands (difficulty). Notably, If task difficulty is viewed as a proxy for the learning stage, these findings point to a strategic shift: early in learning (or under high demands), participants depend on the maintenance period to reorganize information, whereas later in learning (or under low demands), they can apply well-established encoding strategies directly and efficiently during stimulus presentation. This progression closely mirrors skill acquisition, in which novices require extensive post-hoc rehearsal while experts implement efficient routines online.

Unlike traditional neural replay paradigms using explicitly ordered sequences (Foster & Wilson, 2006; Huang et al., 2018; Liu et al., 2019; Liu, Mattar, et al., 2021; Schuck & Niv, 2019), our design presented all items simultaneously, requiring participants to self-organize a sampling order. Despite this, eye trajectories consistently followed a near-optimal path through feature space (e.g., low-to-high frequency), indicating internally generated organization. This aligns with evidence from decision-making and visual search tasks (Hoppe & Rothkopf, 2019; Yang et al., 2016; Zhu et al., 2022), where gaze trajectories reflect planned, efficiency-driven sampling.

Moreover, the consistent tendency to initiate sequences with low-frequency items may reflect an anchoring strategy. This bias resonates with broader theories of magnitude representation, such as the mental number line (Dehaene, 1996; Dehaene et al., 1993), although our data do not provide direct evidence for a spatial left–right mapping.

Over the past decades, eye-tracking has emerged as a powerful tool for probing memory-related cognition (Hannula et al., 2010; Meister & Buffalo, 2016). One particularly striking insight is how rapidly memory expression is reflected in eye movements. Early work established this link by analyzing summary statistics, such as visual sampling duration and saccade vector distributions, which reliably predict memory performance and hippocampal engagement (Althoff & Cohen, 1999; Hannula & Ranganath, 2009; Henderson et al., 2005; Hoffman et al., 2013; Liu et al., 2017; Olsen et al., 2016; Shen et al., 2016). Later studies used scanpath analysis to further demonstrate that during retrieval, individuals’ gaze patterns precisely reinstated those from the encoding phase. (Johansson et al., 2022; Laeng & Teodorescu, 2002; Sahan et al., 2024; Wynn et al., 2020). Extending this work, our study characterizes the fine-grained temporal architecture of sequential eye movements during working memory. The results show that even without external input, gaze reflects internal sampling mechanisms that actively structure and maintain multiple memory representations.

An alternative account is that low-level visual salience drove the observed sequences. Although saliency models (Itti & Koch, 2001) predict that perceptual features can guide sequential gaze during encoding, two observations argue against this interpretation. First, the structured sequence reliably reappeared during the maintenance period, when no visual input was present, indicating a memory-driven mechanism. Second, the systematic shift of sequential sampling from encoding to maintenance with increasing task difficulty reflects flexible cognitive control rather than passive stimulus-driven responses. Together, these findings suggest that sequential eye movements reflect active memory optimization rather than bottom-up salience.

In conclusion, we show that increasing cognitive demands lead to a shift in when optimal eye-movement strategies are expressed—from online sampling during encoding to offline replay during maintenance. These internally generated, near-optimal trajectories demonstrate that eye movements are not merely passive reflections of visual processing but active manifestations of the internal reorganization processes that support memory optimization. By capturing these dynamics in real time, our study positions gaze as both a window onto and a mechanism for adaptive control, offering a powerful noninvasive marker of cognitive load, memory organization, and internal sampling in complex environments.

## Methods

### Participants

Eighty-nine participants (aged 18–35 years) took part in the study: 48 in Experiment 1 (3-item memory) and 41 in Experiment 2 (4-item memory). They were recruited via the internal database recruitment system of Max Planck Institute for Human Cognitive and Brain Sciences in Leipzig. The study was approved by the local Ethics Committee of Leipzig University, Germany (Ethics Approval: Nr 018/23-ek). All the participants were naïve to the purpose of the experiments and gave written informed consent before the experiment and were compensated for their participation with 13 Euros per hour. All participants had normal or corrected-to-normal vision, with no history of psychiatric or neurological disorders. Due to poor eye-tracking data quality, two participants were excluded from Experiment 1 and one from Experiment 2.

### Stimuli and tasks

Visual stimuli consisted of either three (Experiment 1) or four (Experiment 2) items presented simultaneously at different locations (7° eccentricity). Each item comprised two integrated components: (1) an internal grating with one of six possible spatial frequencies (ranging from 0.5° to 8° visual angle, logarithmically spaced), and (2) an outer ring in one of six colors (evenly spaced from blue to green in Commission Internationale de l’Eclairage (CIE) LAB color space). A systematic relationship was established between stimulus features: lower spatial frequencies were consistently paired with greener hues. This created an implicit feature-space structure where color and frequency varied coherently along a single dimension (greener pairing lower frequency; bluer pairing higher frequency), enabling optimal information organization through sequential sampling.

#### Experiment 1 (3-item memory)

Each trial began with a central fixation requirement (gaze within 1° for at least 10 ms) to establish a neutral baseline. Three gratings, varying in spatial frequency and color, then appeared simultaneously for 4 s at 7° eccentricity, arranged in an equilateral triangle configuration for encoding. After stimulus offset, participants re-fixate centrally to initiate the 2 s maintenance phase, where three placeholder circles appeared at original stimulus locations. Another central fixation triggered the retro-cue period, where the three circles reappeared with one filled for 100 ms to direct attention to that specific location, followed by another 2 s display of all circles. Finally, a test stimulus (grating or color ring) appeared at the center alongside the three circles. Participants were required to indicate which memory item matched the test stimulus by clicking the corresponding location, or to press the “C” key if none of the memory items matched.

Therefore, the chance level of memory performance is 0.25. Note that three fixation periods were used to control for spatial biases during memory encoding/maintenance/recalling period. The experiment comprised 216 trials (6 blocks of ∼12 min each) with inter-block eye-tracker calibration, totaling ∼100 minutes (including breaks). As shown in Figure 1B, there were six distinct spatial configurations based on relative stimulus values, with 36 trials for each configuration.

#### Experiment 2 (4-item memory)

Experiment 2 employed the same stimuli and design as Experiment 1, with one critical modification: four items (instead of three) were simultaneously presented for 5 s at 7° eccentricity, positioned at the vertices of an equilateral rectangle, each possessing distinct spatial frequencies and colors. Following stimulus offset, participants maintained the four items in memory for 2.5 s. To account for the greater cognitive load of the 4-item task compared to the 3-item version, we extended both the encoding duration (from 4 s to 5 s) and maintenance period (from 2 s to 2.5 s) to ensure adequate information processing. During the response phase, participants indicated either the location matching the test stimulus or pressed “C” for novel items, resulting in a chance performance level of 0.2. Participants also needed to complete 216 trials (6 blocks of ∼13 min each) with inter-block eye-tracker calibration. In total, it took around 110 minutes (including breaks). From twenty-four possible spatial configurations, we selected twelve for testing (18 trials per configuration) based on signal-to-noise ratio considerations.

### Eye-tracking recordings

Participants sat 60 cm from a monitor (1280 × 1024 pixels, 60 Hz refresh rate) with their head stabilized using a chin rest. An eye tracker (EyeLink 1000 plus, SR Research) was positioned on the table approximately 15 cm in front of the monitor. During the task, gaze was continuously tracked for both eyes simultaneously, at a sampling rate of 1,000 Hz. Before acquisition, we calibrated the eye tracker using the built-in calibration and validation protocols from the EyeLink software.

### Eye-tracking preprocessing

Raw eye-tracking data were converted from EDF to MAT format using the Edf2Mat toolbox with default settings. For each trial, we extracted three epochs of interest: (1) the stimulus presentation epoch (−10 to 4000 ms from stimulus onset), (2) the memory maintenance epoch (−10 to 2000 ms from placeholder onset), and (3) the memory retrieval epoch (−10 to 2100 ms from retro-cue onset). Baseline correction was applied using the pre-event interval (−10 to 0 ms). Blinks were detected by identifying NaN clusters (≥9 consecutive missing samples) in each epoch. To capture pre/post-blink distortions, we extended each blink window by 100 ms on both sides. Gaze data from both eyes were then averaged, producing horizontal (x) and vertical (y) gaze time series per epoch. For data quality control, epochs with more than 50% invalid samples were discarded. Remaining blink artifacts were interpolated using MATLAB’s makima method (a smooth cubic Hermite interpolation). Finally, trials where gaze deviated >10° visual angle from central fixation were excluded to ensure proper gaze. Following this cleaning procedure, the average number of valid trials retained was 198 (SD = 23) in Experiment 1 and 186 (SD = 26) in Experiment 2.

### Eye-tracking analysis

#### Correlation based on the spatial configuration

Since eye movements directly reflect overt attention to specific stimuli, we quantified the strength of item representation by calculating the Euclidean distance between the gaze position and each item’s location. Smaller distances indicated stronger attentional representation of that item. This produced three gaze-to-item distance time courses per epoch (one per item). We hypothesized that eye-movement trajectories would reflect the spatial configuration of items, defined by the specific arrangement of relative feature values (e.g., low, medium, high) across the three fixed stimulus locations (As illustrated in Figure 1B, this resulted in six possible spatial configurations). For example, if a low-frequency item appeared at the top-left, medium at the top-right, and high at the bottom-center, gaze patterns should resemble those of other trials with the same configuration more than those with a different one (e.g., high at top-left, medium at top-right, low at bottom-center). To test this, we computed pairwise correlations of the gaze-to-item distance time courses across all trial pairs. Correlations were then grouped by whether the trial pair shared the same spatial configuration or not. Higher correlations within same-configuration pairs compared to between-configuration pairs would indicate that gaze trajectories reflect stable, value-guided spatial sampling patterns. Note that we averaged the configuration-based correlation values across all items to obtain a single pattern-consistency score per trial pair. This analysis was performed separately for the encoding and maintenance periods. To assess behavioral relevance, we then tested whether individual differences in this pattern-consistency score (i.e., the difference between same– and different-configuration correlations) predicted memory accuracy. The same analysis was repeated in Experiment 2, with the only difference being that it was applied to four items with twelve different configurations.

#### Sequence analysis on memory encoding and maintaining period

To uncover the sequential structure in the eye-movement trajectories, we applied sequence analysis (Liu, Dolan, et al., 2021; Liu, Mattar, et al., 2021) to the gaze-to-item distance time courses, where items were labeled according to their relative frequency values within each trial (item 1: lowest frequency/greenest; item 2: middle frequency/ intermediate hue; item 3: highest frequency/least green). This analysis proceeded in several steps. First, we computed the cross-correlations between each pair of items’ gaze-distance time series across a range of temporal lags (0 to 2000 ms in experiment 1, 0 to 2500 ms in experiment 2), generating an empirical correlation matrix at each lag. Using TDLM framework (Liu, Dolan, et al., 2021), we then modeled these empirical correlation matrices using theoretically-derived transition matrices.

Specifically, we tested two competing sequences: (1) a value-increasing sequence (item 1→item 2→item 3), meaning that the representation of item 1 at time T is expected to predict the representation of item 2’s representation at T+Δt, which in turn predicts item 3 at T+2Δt; and (2) its reverse sequence (item 3→item 2→item 1). To quantify how well each theoretical transition pattern accounted for the observed gaze dynamics, we computed a second-level correlation between the predicted transition matrix and the empirical cross-correlation matrices across lags. All analyses were performed separately for the encoding and maintenance periods.

To assess statistical significance, we conducted a permutation test by randomly shuffling the item labels within each trial and repeating the sequence analysis on these surrogate datasets. This procedure was repeated 200 times, yielding a null distribution of sequence strength values across temporal lags. A significance threshold (p < 0.05) was then determined at each lag based on this null distribution. To correct for multiple comparisons across lags, we used the maximum value approach by setting the highest threshold across all lags as the corrected significance cutoff.

The same analysis was applied to Experiment 2, with adjustments to accommodate the four-item design. Specifically, the empirical correlation matrices were expanded from 3 × 3 to 4 × 4, and the transition matrices were updated accordingly to reflect sequences among four items.

#### Sequential saccades after retro-cue

Saccades were detected using a velocity-based algorithm (Engbert & Kliegl, 2003; Holmqvist et al., 2011), in which gaze-position data were first transformed into a velocity time series by calculating the Euclidean distance between consecutive samples. Saccade onset and offset were identified when gaze velocity exceeded a threshold of 30° of visual angle per second, with the additional constraints that saccade duration had to fall between 10 and 300 ms and that inter-saccadic intervals were at least 20 ms to ensure distinct, physiologically plausible events.

To examine whether the retro-cue directed gaze toward the cued location, we analyzed the landing position of the first saccade after cue onset, specifically testing whether it was closer to the cued position than to the uncued locations. In trials where the first saccade was directed to the cued location, we further analyzed the subsequent (second) saccade to assess whether it followed an optimal mental scanning trajectory based on item value order (i.e., item 1→item 2→item 3). For example, if the first saccade landed on item 1 (lowest frequency/greenest), the second saccade was expected to target item 2 (middle frequency), rather than item 3 (highest frequency). Similarly, if the first saccade landed on item 2, the second saccade was more likely to shift toward item 3 than item 1, consistent with a value-based sampling sequence. To test this, trials were selected based on the following criteria: (1) the first saccade was directed to the cued location, (2) the cued item was either item 1 or item 2, and (3) at least two saccades occurred within the trial. Participants with fewer than 20 valid trials based on these criteria (N = 4) were excluded from the analysis. For the remaining trials, we compared the forward, value-consistent transitions (item 1→item 2; item 2→ item3) with alternative transitions (item 1→item 3; item 2→item 1), to determine whether the second saccade reflected a structured, optimal sampling strategy.

### Author contributions

Q.H. and C.F.D. conceived the initial idea. Q.H. designed the experiment and developed the tasks. Q.H. acquired the data. Q.H. planned and performed analyses. Q.H. and C.F.D. discussed the results. Q.H. wrote the initial draft. Q.H. and C.F.D. finalized the manuscript.

## Acknowledgments

This work was supported by Humboldt Research Fellowship for Postdocs to Q.H. and Max Planck Society. We thank Nicholas Menghi, Simone Vigano, Yangwen Xu, and Marit Petzka for their helpful comments.

## Supplementary figure

**Supplementary figure 1.**
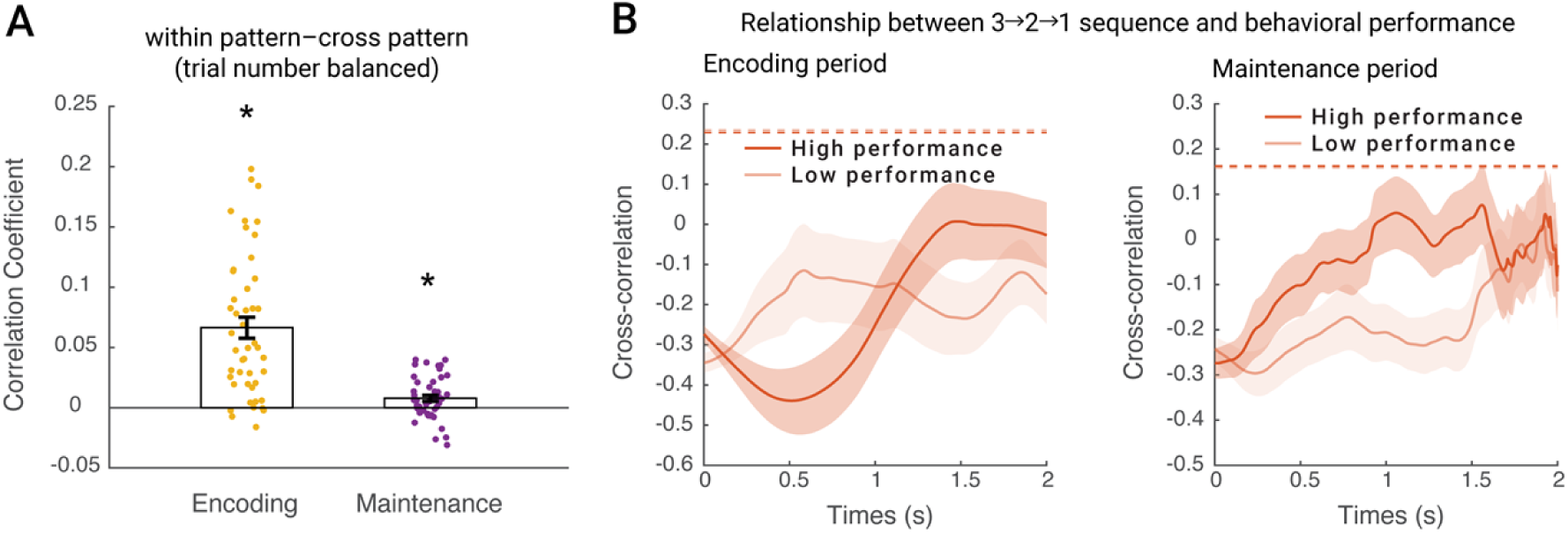
(**A**) To account for unequal trial numbers between the within-pattern and cross-pattern conditions in Experiment 1, we randomly subsampled the cross-pattern trials 100 times and averaged the results across these iterations. The difference in pattern similarity (within-pattern minus cross-pattern) was then computed separately for the encoding (yellow) and maintenance (purple) periods. *: p < 0.05. **(B)** To assess the behavioral relevance of the 3→2→1 sequence, we compared sequence strength between high (dark red) and low-performance (light red) groups, based on a median split of memory accuracy (N = 23 per group). Results are shown separately for the encoding period (left panel) and the maintenance period (right panel).

**Supplementary figure 2.**
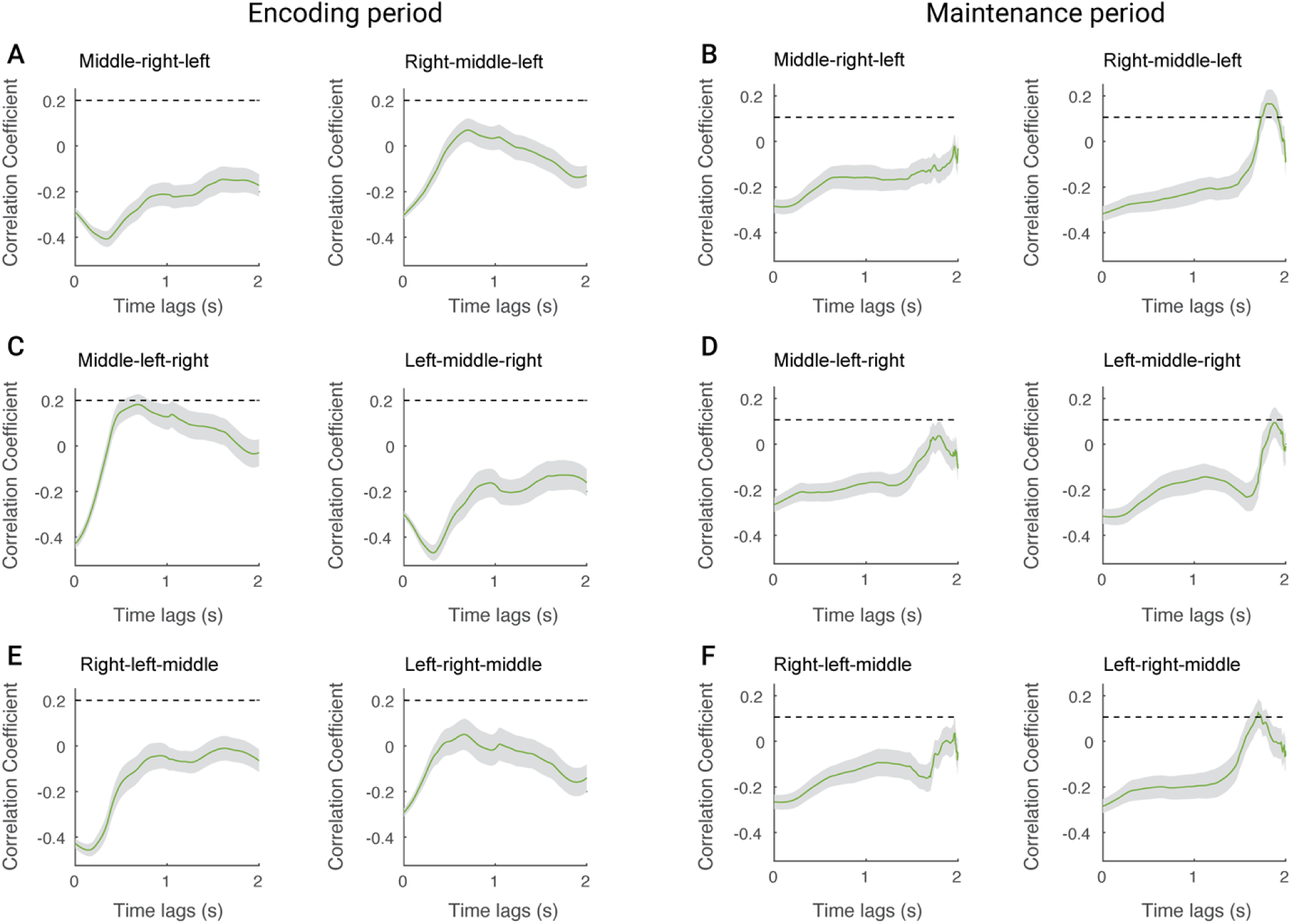
Six possible sequences are based solely on spatial locations. To rule out the possibility that sequential eye-movement patterns were driven by systematic spatial biases rather than value-based optimal organization, we relabeled items based only on their spatial positions and re-ran the same sequence analysis between them during encoding (panels A–C) and maintenance (panels D–F). No significant sequential patterns were observed, confirming that the original effects were not explained by spatial location alone.

**Supplementary figure 3.**
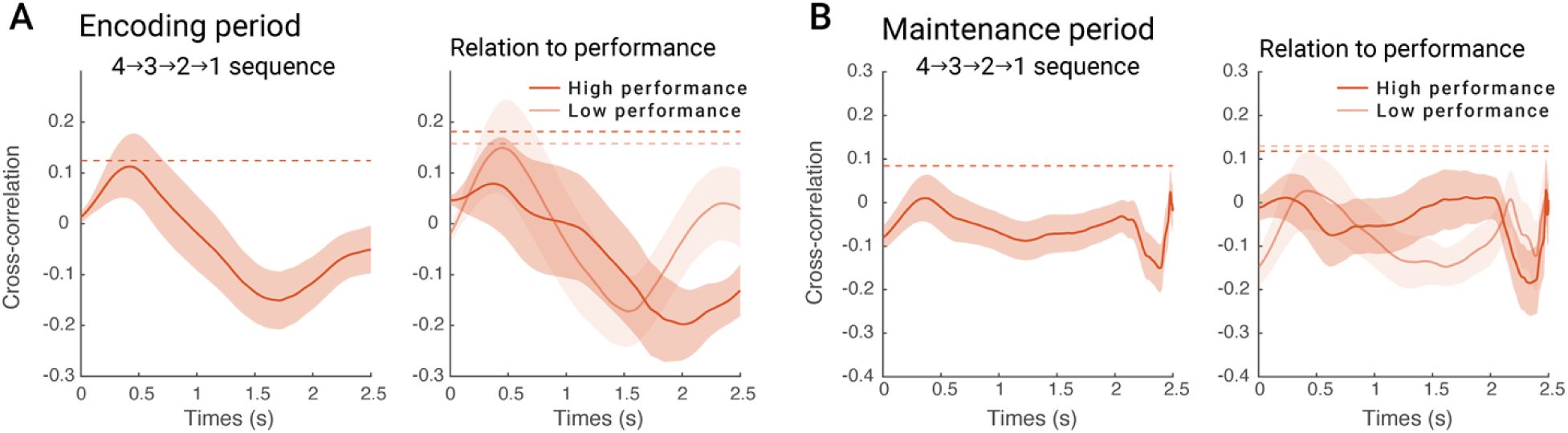
Sequence strength of the 4→3→2→1 pattern in Experiment 2. The reverse value-based sequence (4→3→2→1) was not significant during either the encoding period (**A–Left**) or the maintenance period (**B–Left**). To examine potential individual differences, participants were divided into high (dark red) and low-performance (light red) groups based on a median split of memory accuracy. However, no significant sequence was observed in either group during encoding (**A–Right**) or maintenance (**B–Right**), suggesting that the 4→3→2→1 sequence does not contribute meaningfully to memory performance.

